# Evidence for Toxin-Encoding Coinfections Driving Intransitive Dynamics Between Allelopathic Phenotypes in Natural Yeast Populations

**DOI:** 10.1101/2025.07.30.667734

**Authors:** Tommy J. Travers-Cook, Emmy Gonzalez-Gonzalez, Jukka Jokela, Kayla C. King, Sarah Knight, Claudia C. Buser

## Abstract

Competitive intransitivity — non-hierarchical interactions, such as those exemplified by the rock-paper-scissors game, where no single competitor wins outright — has been proposed as a key mechanism for maintaining biodiversity; however, empirical evidence supporting the importance of intransitivity remains limited. Natural populations of *Saccharomyces cerevisiae* often include strains harboring totivirus-satellite coinfections that encode a lethal toxic glycoprotein capable of eliminating competing yeast strains. Killer strains are sparsely distributed in natural populations, despite their competitive advantage. Yeast isolates occasionally exhibit toxin resistance, but it remains untested whether they can outcompete and replace killer strains. Similarly, the persistence of toxin-susceptible yeast is not well understood —particularly whether they can invade resistant populations in the absence of killers, thereby completing an intransitive loop. In a multi-year collection of yeast isolates from vineyards across New Zealand, we observed a near-complete disappearance of a previously common killer yeast genotype of *S. cerevisiae* over consecutive years. Using space-time-shift competition assays, we demonstrate that strains sympatric to this killer genotype were ubiquitously resistant, unlike the allopatric strains that were frequently eliminated in competition assays. Furthermore, the extinction of the focal killer genotype appears to have enabled the emergence of toxin-susceptible competitors in sites formerly occupied by the killer genotype. Our findings suggest that the competitive advantage of toxin production is evident in natural populations but appears to be eroded when resistance evolves in competitors of the focal killer genotype. We suggest that such killer-resistant-susceptible polymorphisms are being maintained by evolutionary dynamics akin to rock-paper-scissors-*like* intransitivity, driven by the invasion of susceptible strains after costly resistance has driven killer strains to extinction in natural populations, all being driven by toxin-encoding coinfections.

## Introduction

Competitive intransitivity refers to interactions among competitors that are not strictly hierarchical but instead form loops in which no single competitor is universally dominant. Rock-paper-scissors (RPS) exemplifies an intransitive interaction, where winners emerge in pairwise match-ups, yet no overall winner exists within the intransitive loop. The role of intransitivity in promoting species coexistence and maintaining population-level polymorphism is of considerable theoretical interest (May & Leonard 1975; Laird & Schamp 2006; Reichenbach et al. 2007; Alcántara, Pulgar & Rey 2017), and has also been documented in various natural systems (Sinervo & Lively 1996; Soliveres et al. 2015; Ulrich et al. 2016; Vandermeer & Perfecto 2023).

Intransitivity has been proposed as a mechanism for promoting coexistence even in the absence of niche differentiation (Soliveres & Allan, 2018; Allesina & Levine, 2011; Laird & Schamp, 2006). Similarly, spatial structure in competitively homogenous environments that restricts mixing can facilitate coexistence by promoting local rather than global interactions (Amarasekare 2003). In populations with low niche differentiation, intraspecific competition tends to be reduced under low-density conditions. However, when individuals aggregate in high-density niches, this may generate a patch-like population structure characterised by local interactions among subsets of competitive polymorphs, creating conditions under which intransitive dynamics can arise and be maintained.

There has been growing interest in the role of intransitivity in maintaining polymorphism and species diversity in microbial populations and communities, respectively (Verdú, Alcántara & Navarro-Cano, 2023). In microbial systems, interference competition typically involves antagonistic allelopathy—such as the production of toxins—and often results in a clear competitive outcome, with the winner being the organism that successfully eliminates its competitor (Chao & Levin, 1981; Riley & Gordon, 1999; Ghoul & Mitri, 2016). Natural metapopulations of microbial species commonly maintain polymorphism in toxin production, resistance to toxins and apparent susceptibility to toxins. Maintenance of such polymorphism requires rapid evolution of toxin resistance during ecologically relevant time scales. However, resistance is not ubiquitous. In some environments, selection may favor susceptibility—particularly when resistance carries a cost—which could make these populations vulnerable to invasion by strains that produce allelopathic toxins, completing an intransitivity loop.

The budding yeast *Saccharomyces cerevisiae* is typically scarce but consistently present across diverse habitats in temperate and tropical climates (Wang et al., 2012). However, it dominates sugar-rich substrates due to its superior fermentation capacities, a result of its Crabtree-positive nature (Merico et al., 2007). Some strains of *S. cerevisiae* produce allelopathic toxic glycoproteins that are often found to be lethal for intra- and interspecific competitors (Boynton, 2019; Magliani et al., 1997; Schmitt & Breinig, 2002; Starmer et al., 1987). Production of these allelopathic toxins is encoded either by a totivirus-satellite coinfection, or by chromosomal genes (Schmitt & Breinig, 2002, 2006). Totiviruses are monopartite dsRNA viruses found in species across the Dikarya subkingdom of fungi (Liu et al., 2012), that tend to possess two overlapping open reading frames (ORF) encoding a capsid protein and an RNA-dependent RNA polymerase for replication (Dinman et al., 1991; Dinman & Wickner, 1994; Fujimura et al., 1992; Icho & Wickner, 1989). Satellites instead encode a pre-processed toxin on their single ORF (Dignard et al., 1991; Hanes et al., 1986; Schmitt & Tipper, 1995) and rely on totivirus gene products for persistence in the host cell (Bostian et al., 1980). Satellite-encoded toxin production acts analogously to toxin-antitoxin systems in bacteria (Jurenas et al., 2022) because totivirus satellites also encode immunity (Dignard et al., 1991; Hanes et al., 1986). Loss of immunity through loss of the satellite results in susceptibility to toxins produced by the clonal neighbours that maintain the coinfection (Travers-Cook et al., 2023). Competing strains that are not coinfected by the totivirus-satellite combination are susceptible to killer toxins unless they have independent mutations providing resistance (Bostian et al., 1984; Breinig et al., 2002; Delapena et al., 1981; Magliani et al., 1997; Marquina et al., 2002; Riffer et al., 2002).

Allelopathic toxins select for resistance in strains with toxin-susceptible genetic backgrounds (Pieczynska et al., 2016). Toxins eliminate competitors most effectively when the killer strain is in high density (Greig & Travisano, 2008), which suggests that the ecological advantage of a killer phenotype is to suppress invading strains from the resource patch (Travers-Cook et al., 2023). Despite the competitive advantage of the killer phenotype, strains that produce toxins are not statistically overrepresented in *S. cerevisiae* populations and collections (Chang et al., 2015; Crabtree et al., 2023; Travers Cook et al., 2025). Coexistence with a killer strain should require resistance to killer toxins. Where killer strains have disappeared through the evolution of resistance, dispersal may enable the influx of toxin-susceptible strains into resistant habitat patches (Boynton, 2019). If resistance bears a cost, toxin-susceptible strains may be able to replace resistant strain patches where killer strains have gone extinct (Boynton, 2019). Under such a scenario of toxin- and resistance-associated costs, the distribution and turnover of competitive polymorphism may be related to these discrete and unambiguous phenotypic differences with regards to toxin production, forming an killer-sensitive-resistance intransitive loop analogous to rock-paper-scissors.

In this study, we ask whether toxin-encoding totivirus-satellite coinfections drive rock-paper-scissors-like intransitive dynamics in *Saccharomyces cerevisiae* populations. We hypothesise that killer strains replace toxin-susceptible strains, as evidenced by the absence of toxin-susceptibility among strains coexisting with killers. Conversely, we hypothesise that toxin-susceptible yeast can invade resistant patches, as evidenced by their presence in sites formerly occupied by killers but now dominated by resistant genotypes. In a collection of vineyard-associated *S. cerevisiae* yeast isolates from New Zealand, we identified a yeast genotype that was associated with toxin-encoding totivirus-satellite coinfections across the whole study region. This killer strain was found to have a patchy distribution across vineyards sampled in 2018, and experienced a near-complete disappearance in the following year. Using a space-time shift design, we competed this killer genotype against sympatric and allopatric competitors from years where the killer genotype was present and absent. We found that all sympatric contemporaries of the killer genotype were toxin-resistant, unlike in allopatry, demonstrating that the satellite-encoded toxins effectively select for resistance in sympatric strains. We also found that in the absence of the killer genotype, susceptible strains re-emerged. Our study demonstrates that the presence of toxin producing genotypes determines the spatiotemporal distributions of toxin sensitivity and maintains polymorphism through evolutionary dynamics described by rock-paper-scissors type intransitive polymorphism.

## Materials and Methods

### Yeast Isolate Collection, Genotyping and Infection Screening

The yeast isolates used in the study are the same as described in Travers-Cook et al. (2025). To briefly summarise, 536 *S. cerevisiae* isolates from 25 vineyards over three years (2018, 2019 and 2021) were sampled from the Hawke’s Bay and Marlborough regions of New Zealand. All isolates were genotyped with eight microsatellite loci and infection states was determined by dsRNA extractions (Crabtree et al., 2019; Richards et al., 2009).

### Analysis to Identify a Killer Genotype with a totivirus-satellite coinfection

Using the genotype and infection status data, we tested for significant changes in the prevalence of genotypes with the toxin-encoding satellites using a non-parametric Wilcoxon signed-rank test (R Core Team, 2023). We found a significant decline in the presence of a genotype that consistently hosted the toxin-encoding totivirus-satellite coinfections (Wilcoxon signed rank test: V = 88.5, *p* = 0.003; Figure 1). This genotype, hereafter referred to as the killer genotype, was found to dominate many vineyards in 2018, but was nearly-completely absent from the 2019 collections of the following year, and had experienced a complete local extinction from most vineyards that it dominated in the previous year. Across the vineyards of 2018, the killer genotype was found to vary from dominance to complete absence in others. This spatiotemporal variation in the prevalence of a killer genotype enabled us to explore how the presence of a killer phenotype influences the distribution of toxin resistant and toxin susceptible strains.

**Figure 1.**
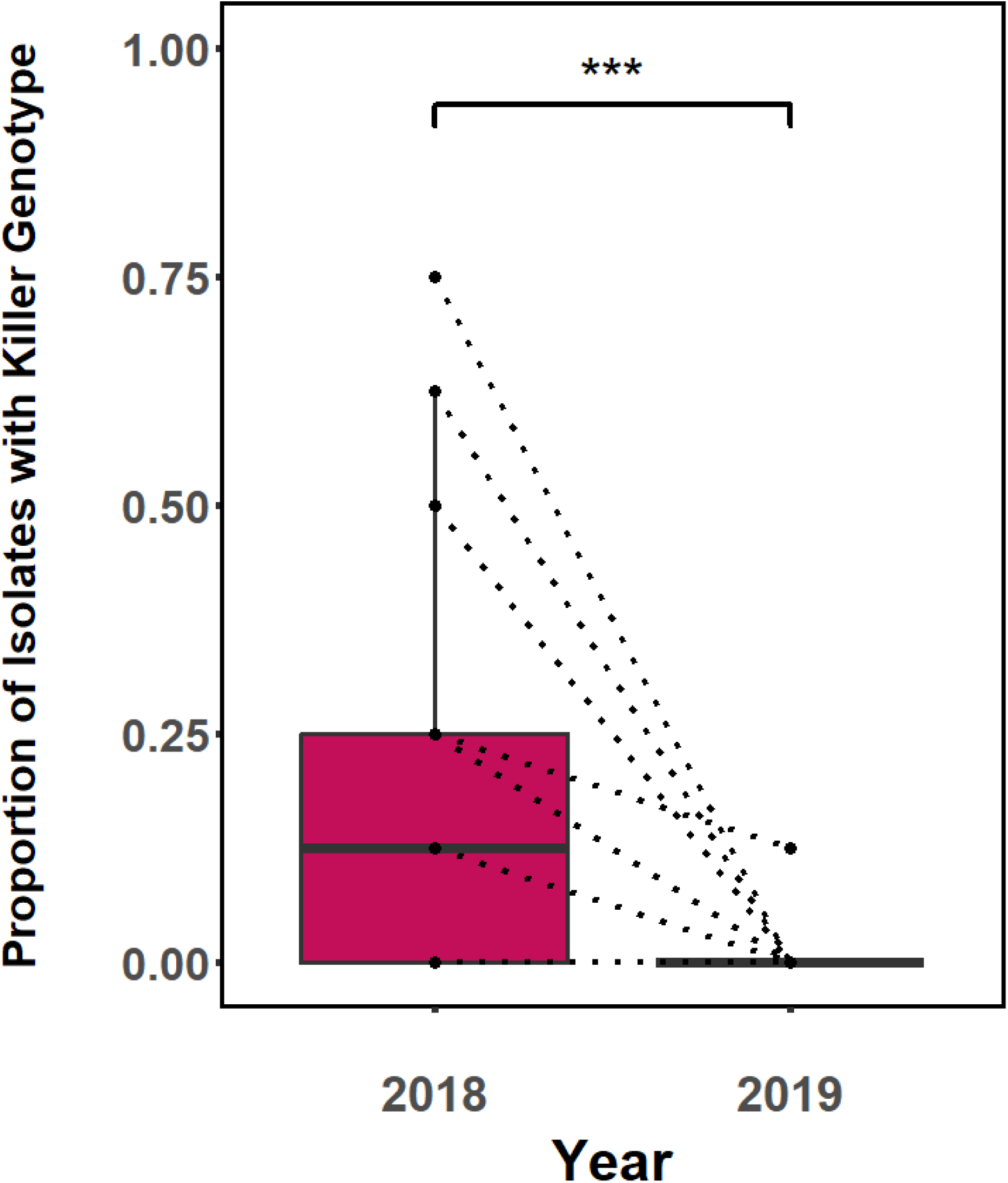
Spatiotemporal variation in the prevalence of the common killer genotype in vineyards of New Zealand. Individual points are vineyards. Dotted lines connect vineyard sites across years.

### Space-Time Shift Experimental Design and Yeast Isolate Selection

Using this common killer genotype of 2018, we tested whether the prevalence of resistance among non-killers was dependent on whether these strains were of sympatric or allopatric origin. To do so, we used a space-time shift design, by competing isolates of the killer genotype against strains from the same and different locations across the years where it was present and absent (Figure 2). We labelled these strains, indicating whether they were sampled from a vineyard where the common killer genotype was observed or absent from, as “*Sympatry*” and “*Allopatry”*, respectively. For the experiment we knew if the killer phenotype was present in 2018 and 2019. We labelled the time-factor as “*Contemporary*” when the strain was present in 2018 (the year the killer genotype was widespread), and “*Future*”, if the strain was sampled in 2019 when the killer genotype was no longer present. We used a full factorial design, giving us four treatment combinations: *“Contemporary x Allopatric”, “Contemporary x Sympatric”, “Future x Allopatric” and “Future x Sympatric”*. We randomly sampled 34-36 combinations of isolates of the killer genotype and of competitor genotypes, from each of the four treatments, without replacement, to test for differences in resistance (Table S1; R Core Team, 2023). Though the focal killer genotype was found to be rare in a number of vineyards, for competition assays we only chose isolates of the genotype from vineyards where the genotype dominated (occupancy of ≥50% of more). We took from sites where the focal killer genotype was dominant because killer effectiveness is positively density dependent (Greig & Travisano, 2008). Yeast from allopatric sites were however from vineyards where the killer genotype was entirely absent, as this would minimise any potential effect of the killer being present but undetected. All of the competitors of the killer genotype were verified as belonging to different genotypes to the killer genotype.

**Figure 2.**
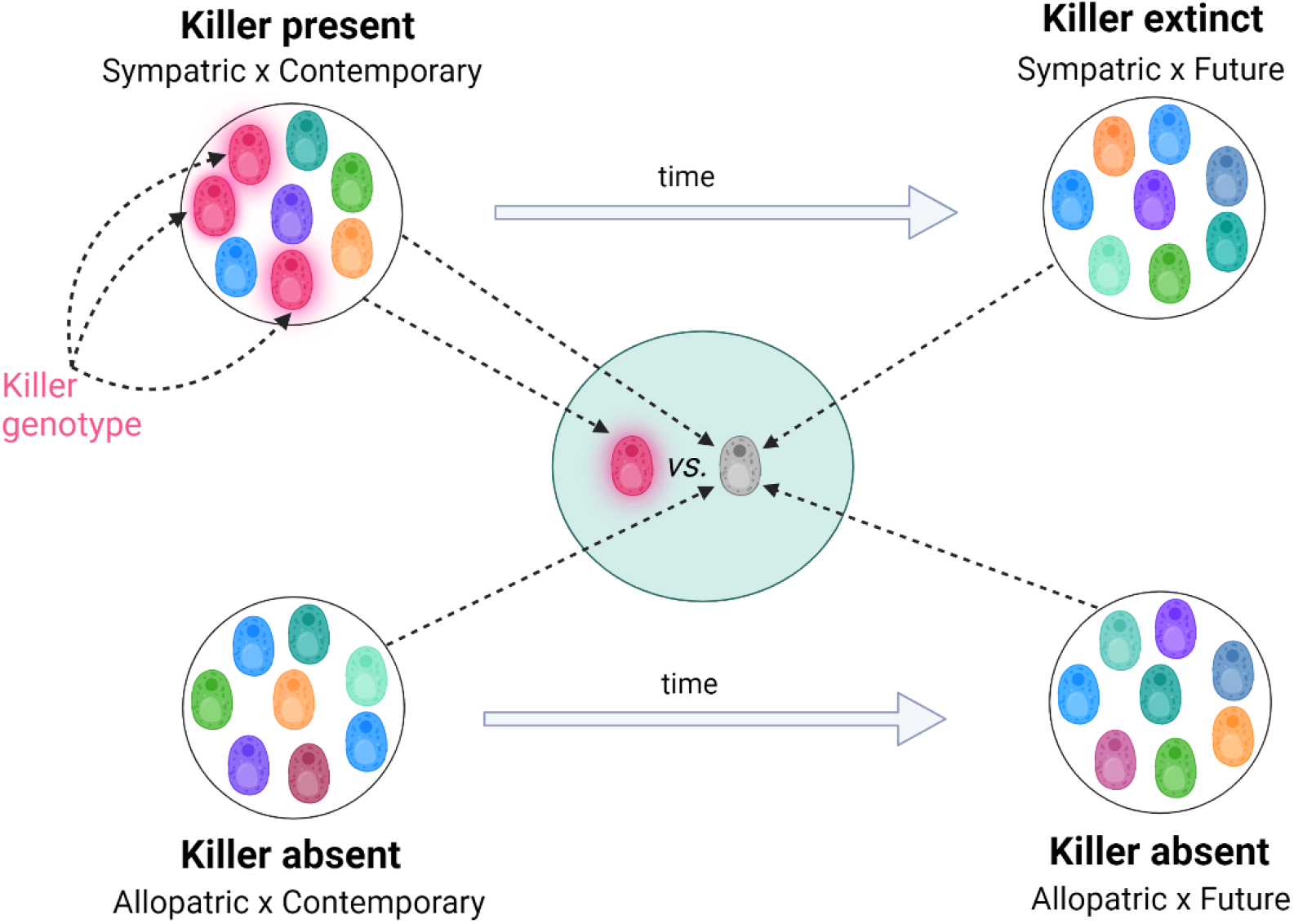
Experimental design for the space-time shift experiment. Yeast isolates of a specific genotype, which exhibit the killer phenotype, were designated as the “killer genotype”. Yeast isolates were then randomly sampled from a collection, from sites where the killer genotype was present (“*Sympatry*”) and sites where it was always absent (“*Allopatry*”). The competitors of the focal killer genotype were taken from two different years: 2018, when the killer genotype was present (“*Contemporary*”), and 2019, when the killer genotype had undergone extinction (“*Future*”). The experimental design followed a full factorial approach with vineyard (*Sympatry vs. Allopatry*) and year (*Contemporary x Future*) as treatments, resulting in four distinct levels: *Sympatric x Contemporary* (where the killer genotype was present), *Allopatric x Contemporary* (where the killer genotype was absent in the year that it was present), *Sympatric x Future* (where the killer genotype was historically present but extinct in 2019) *and Allopatric x Future* (where the killer genotype was always absent). The *Allopatric* conditions across both years were treated as controls to establish a baseline level of resistance in vineyards where the killer genotype had never been observed.

### Space-Time Shift Inhibition Assays

All yeast isolates were stored in 15% glycerol at −20 °C in 96 well plates prior to reanimation for the experiments. Yeast isolates were grown in standard liquid YPD medium (1% yeast extract, 2% glucose, 2% peptone) prior to their use in the experiments. For testing the killing abilities of yeast throughout, pH-adjusted (pH = 4.5) YPD solidified with 2% agar and supplied with 0.4% methylene blue (MB) for dead cell staining was used. Competitors of the focal genotypes were prepared by inoculating 25 mL of liquid YPD media with 20 µL of yeast isolate glycerol stock, before incubating for 22 hours at 30 °C with a constant shake of 200 rpm. These cultures were then diluted 1:100 with YPD media. 200 µL of each dilution was then spread evenly across the previously described pH-adjusted, agar-supplied YPD-MB petri dishes. In individual wells of a 96 well plate, 190 µL of YPD media was inoculated with 10 µL of each isolate with the focal genotype assignment and incubated for 24 hours at 30 °C. Once grown, yeast cells were concentrated by centrifugation (3000 x *g* for 2 min) and removal of the top 100 µL of YPD from each well without disturbing the yeast pellet. After thorough resuspending of yeast cells for each isolate, 1.5 µL of each was pipetted onto the corresponding competitor lawn. Plates were then incubated at 25°C for 96 hours. Plates were then checked for methylene blue halos indicating growth inhibition of the lawn strain (Lopes & Sangorrín, 2010).

### Data Analysis

Each killer assay was classed as a binary response variable, either as *halo present* (1 – toxin-susceptible) or *halo absent* (0 – toxin-resistant). To establish whether the main effects of cohabitation (“*Allopatric*” or “*Sympatric*” competitor of killer genotype) and concurrence (“*Contemporary*” or “*Future*” competitor of killer genotype), as well as their interaction, affect the binary outcome of resistance-sensitivity to the killer genotype’s toxins, we began with a GLMM framework with the vineyards of origin as random effect. We found that virtually none of the variance could be explained by the vineyards from which the killer genotype and their competitors came from. This, in combination with inflated standard errors due to perfect separation in one of the factor levels, justified employment of Firth’s penalised likelihood logistic regression using the logistf package in R (Heinze & Schemper, 2002; Heinze et al., 2013, Puhr et al., 2017; R Core Team, 2023). Firth’s penalised likelihood was used because it could address the issue of complete (perfect) separation by adding a bias-reducing penalty to the likelihood function.

## Results

Using Firth’s penalised likelihood logistic regression, we found a significant interaction effect of concurrence and cohabitation on the prevalence of resistance amongst competitors of the focal killer genotype (Table 1). In the absence of toxin producers (*Allopatry*), toxin-susceptible and toxin-resistant competitors were close to equally present (Figure 3). Concurrence (*Contemporary* versus *Future*) was not found to have a significant effect on prevalence of resistance to the focal killer genotype’s toxins, seemingly due to the consistencies of resistance levels in allopatric populations across time. Amongst sympatric contemporaries of the focal killer genotype, resistance to toxin production was found to be universally present (*Sympatric* x *Contemporary*; Figure 3), which contributed considerably to a significant effect of cohabitation (Table 1). In the absence of the focal killer genotype in 2019, toxin-sensitivity was found to return to sites previously occupied by toxin producers (*Sympatric x Future*), demonstrating an interaction between cohabitation and concurrence. Whilst resistance levels declined in 2019 after the killer genotype went extinct (*Sympatric x Future*), the prevalence of toxin sensitivity in these vineyards did not increase to levels observed in allopatric sites across both years (*Allopatric x Contemporary* and *Allopatric x Future*).

**Table 1.** Effects of cohabitation (*Allopatry* versus *Sympatry)*, concurrence (*Contemporary* versus *Future*) and their interaction on resistance to the killer genotype’s toxins, using Firth’s penalised likelihood logistic regression.

| Variable | Coefficient | SE | $\chi^2$ | P |
| --- | --- | --- | --- | --- |
| *Cohabitation* | 4.073 | 1.462 | 23.608 | <0.001 |
| *Concurrence* | 0.109 | 0.467 | 0.054 | 0.816 |
| *Cohabitation x Concurrence* | 3.119 | 1.545 | 7.484 | 0.006 |

| Variable | Coefficient | SE | $\chi^2$ | *P* |
| --- | --- | --- | --- | --- |
| *Cohabitation* | 4.073 | 1.462 | 23.608 | <0.001 |
| *Concurrence* | 0.109 | 0.467 | 0.054 | 0.816 |
| *Cohabitation x Concurrence* | 3.119 | 1.545 | 7.484 | 0.006 |

**Figure 3.**
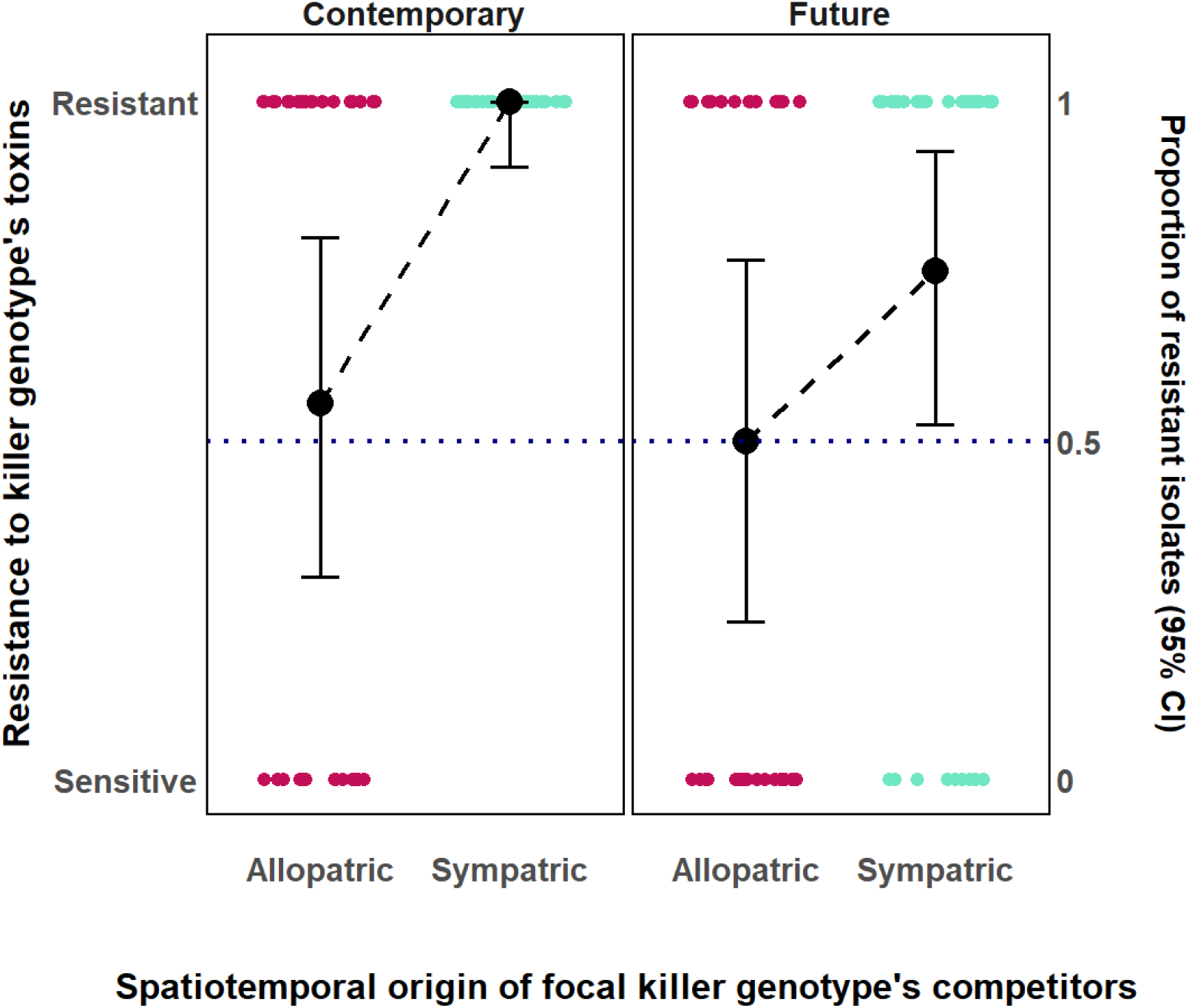
The prevalence of resistance to the satellite-encoded toxins of the killer genotype. The *x*-axis shows whether competitor isolates were sampled from the same site (*Sympatric*) or a different site (*Allopatric*) in relation to the focal killer genotype, and the facets show whether these isolates were from the year the killer genotype was present (*Contemporary*), or the year that it was largely absent (*Future*). The y-axis has a binary scale of resistance-sensitivity to the toxins production by the focal MLG. Individual points are individual assay results. Large black dots represent the proportion of assays that are toxin-susceptible and are skewed towards the response level (susceptible or resistant) that is most prevalent. The 95% confidence intervals were constructed using the Wilson score interval and displayed as error bars for each treatment.

## Discussion

Competitive intransitivity has been suggested as a key mechanism promoting species coexistence and the maintenance of population-level polymorphisms (Soliveres & Allan 2018). Progress in understanding competitive intransitivity has however been hampered by ambiguities in how to assess pairwise competitive abilities and coexistence (Soliveres and Allan 2018). In microbial systems, where toxin-based interference competition is pervasive (Ghoul & Mitri 2016), the question remains as to whether intransitive loops can sustain the diversity observed. *Saccharomyces cerevisiae* is often coinfected by totiviruses and their satellite nucleic acids, the latter of which encodes toxic glycoproteins capable of eliminating competitors of the yeast host (Schmitt & Breinig, 2002, 2006). While killer yeast can select for resistance under controlled laboratory conditions (Pieczynska et al. 2016), it is not known how they structure natural populations, though it is plausible that the killer-susceptible-resistance phenotypic polymorphism can be maintained by rock-paper-scissor-*like* intransitive dynamics (Boynton 2019; Travers-Cook et al. 2023). Given the clear distinction of competitive outcomes (presence-absence of inhibition) and definability of patches for spatial structure (sympatry vs allopatry) in this system, we tested this by competing a toxin-producing killer yeast genotype against conspecifics sampled from both sympatric and allopatric populations, including time points when the killer genotype was present and later extinct (Figure 1), using a time–space–shift experimental design (Figure 2).

As hypothesised, we found that toxin-resistance emerged in association to previous exposure to the killer genotype (Figure 3), demonstrating with wild isolates what has been observed in lab settings (Pieczynska et al. 2016). Toxin-resistant and -susceptible strains were found in all treatments except amongst the ubiquitously toxin-resistant sympatric strains that concurrently cohabited vineyards alongside the focal killer genotype (*Sympatric x Contemporary*; Figure 3). We can assume that allopatric populations have been relatively free of toxin exposure and perhaps reflect the baseline-level for resistance in patches where the killer genotype was not observed to be present. The prevalence of toxin sensitivity in allopatric treatments (*Allopatric x Contemporary* and *Allopatric x Future*) were approximately 50% (Figure 3). Given that we find toxin susceptibility to be frequent in allopatric contemporaries (*Allopatry x Contemporary*), as well as in the other treatments, the results demonstrate that susceptible strains were being eliminated by the killer genotype in sympatric vineyards prior to the experiment. Our findings on the prevalence of sensitivity in allopatric vineyards for *S. cerevisiae* are consistent with other studies of resistance in *Saccharomyces* (Chang et al. 2015).

This standalone result demonstrates that killer phenotypes are structuring local populations by completely removing toxin susceptible genotypes that would be present if the killer genotype was absent. In the process, resistant competitors have an advantage and resistance is selected for. Resistant strains may also invade vineyard patches where the killer genotype is present post-elimination of susceptible strains. Susceptible strains are likely incapable of invading because toxin production is most effective at high killer strain densities (Greig & Travisano, 2008), and there is an apparent competitive advantage. The pH at which the artificial fermentation of the wine grapes takes place (< pH3.6; Travers Cook et al. 2025) is below the optimal range of pH at which toxin production is activated in *S. cerevisae* (~pH4-5; Greig & Travisano, 2008; McBride et al., 2008). Selection for resistance is occurring prior to fermentation, as evidenced by the low levels of sensitivity observed in sympatric sites even after the killer genotype disappears (Sympatric × Future; Figure 4). If selection were instead occurring during the artificial fermentations, we would not observe this lag effect of the killer phenotype on the subsequent year’s population. Instead, sensitivity levels would be expected to not differ significantly from the allopatric baseline.

At some point between our 2018 (*Contemporary*) and 2019 (*Future*) sampling points, the killer genotype appears to have disappeared from the vineyards where it was common before. This genotype’s near-complete extinction demonstrates that the evolution of resistance was likely driving the genotype to near extinction in synchrony across localities. We can assume that toxin production comes with costs that enable resistant genotypes to replace the killer strain. The costs of toxin production can be observed as a slower growth rate and a longer lag phase relative to strains that do not produce toxins (Greig & Travisano, 2008; Pieczynska et al., 2016; Pintar & Starmer, 2003; Wloch-Salamon et al., 2008). Trade-offs between growth and toxin production are expected, as there are inherent energy and resource costs of making the toxin as well as costs of maintaining the totivirus-satellite coinfections. The totivirus is known, for example, to induce proteostatic stress in *S. cerevisae* (Chau et al., 2023). If evolving and maintaining toxin resistance is energetically cheaper than toxin production, resistant strains may be fitter in sympatry than killer strains, which would lead to an eventual replacement of killers (Boynton, 2019). The hypothesis that toxin production is costlier than resistance is, however, unresolved. More detailed studies are needed to determine how the costs of toxin production and costs of resistance relate to the disappearance of the toxin producing killer genotype.

In the absence of the killer genotype, sensitivity emerged in sympatry with resistance (*Sympatric x Future)*. This suggests that in the absence of the killer genotype, rare susceptible genotypes can invade patches occupied by their resistant counterparts, or that some resistant genotypes mutate to become susceptible. We hypothesize that selection for sensitivity in the absence of the killer genotype results from the costs of resistance, which have yet to be demonstrated. An ecological explanation for temporal polymorphism in the yeast populations would be a metapopulation level random background dispersal of each yeast phenotype into each patch. Dispersal in *S. cerevisiae* populations is largely through fruit flies and other fruit-attending insects (Quan and Eisen 2018; Buser et al. 2014), though human activities are known to contribute (Goddard et al. 2010). In the absence of the killer strain, dispersal would restore susceptible strains to patches that are initially occupied by resistant strains and killer strains in patches that are initially occupied by susceptible strains. Under either circumstance, the reemergence of toxin susceptibility in patches previously occupied by killer genotypes following their extinction supports our hypothesis that killer–susceptible–resistant polymorphism is maintained through an intransitive loop, where susceptibility resurfaces once killers are absent. While rock-paper-scissor-*like* competitive intransitivity has been demonstrated under laboratory conditions (Kerr et al. 2002), evidence in microbial systems using wild isolates has remained elusive. However, the patterns observed in our results align closely with the predictions of an intransitive dynamic.

In the time shift experiment we observed that sensitivity emerged in the absence of a formerly dominant killer genotype, but we did not observe killer strains invade populations. Intransitivity theory would predict that toxin producers can invade patches occupied by susceptible strains but not those where resistance is ubiquitous (Boynton, 2019; Travers-Cook et al., 2023). Intransitive competition does not inherently account for how evolutionary changes in killer genotypes—and selection favoring these novel killer phenotypes—can drive phenotypic transitions in their competitors, such as a shift from resistance to sensitivity (analogous to a player in rock-paper-scissors altering strategies). Consideration of how genotypes may shift from states may be necessary to integrate evolutionary biology into the theory and empirical testing of intransitive competition.

Our findings contribute to the limited but growing evidence for competitive intransitivity in natural systems, while underscoring the value of rock-paper-scissor-*like* loops as a robust and promising model for identifying and quantifying intransitivity (Soliveres & Allan 2018). We present a tractable system for assessing the predictions of rock-paper-scissor-*like* intransitivity in natural populations.

## Supplementary material

All supplementary materials have been submitted with the manuscript for review.

## Data availability

All data and scripts necessary to replicate the results will be made available prior to submission.

## Acknowledgements

The authors would like to thank Soon Lee, for his involvement in collecting the yeast isolates from vineyards in New Zealand (Travers-Cook et al., 2025).

